# Exploring Conformational Transitions and Free Energy Profiles of Proton Coupled Oligopeptide Transporters

**DOI:** 10.1101/651646

**Authors:** Mariana R. B. Batista, Anthony Watts, Antonio J. Costa-Filho

**Affiliations:** Ribeirão Preto School of Philosophy, Sciences and Letters, University of São Paulo - Brazil; Department of Biochemistry, University of Oxford, South Parks Road, Oxford - UK

## Abstract

Proteins involved in peptide uptake and transport belong to the proton-coupled oligopeptide transporter (POT) family. Crystal structures of POT family members reveal a common fold consisting of two domains of six transmembrane *α* helices that come together to form a “V” shaped transporter with a central substrate binding site. Proton coupled oligopeptide transporters operate through an alternate access mechanism, where the membrane transporter undergoes global conformational changes, alternating between inward-facing (IF), outward-facing (OF) and occluded (OC) states. Conformational transitions are promoted by proton and ligand binding, however, due to the absence of crystallographic models of the outward-open state, the role of H^+^ and ligands are still not fully understood. To provide a comprehensive picture of the POT conformational equilibrium, conventional and enhanced sampling molecular dynamics simulations of PepT_*st*_ in the presence or absence of ligand and protonation were performed. Free energy profiles of the conformational variability of PepT_*st*_ were obtained from microseconds of adaptive biasing force (ABF) simulations. Our results reveal that both, proton and ligand, significantly change the conformational free energy landscape. In the absence of ligand and protonation, only transitions involving IF and OC states are allowed. After protonation of the residue Glu300, the wider free energy well for Glu300 protonated PepT_*st*_ indicates a greater conformational variability relative to the apo system, and OF conformations becames accessible. For the Glu300 Holo-PepT_*st*_, the presence of a second free energy minimum suggests that OF conformations are not only accessible, but also, stable. The differences in the free energy profiles demonstrate that transitions toward outward facing conformation occur only after protonation and, probably, this should be the first step in the mechanism of peptide transport. Our extensive ABF simulations provide a fully atomic description of all states of the transport process, offering a model for the alternating access mechanism and how protonation and ligand control the conformational changes.

**Graphical TOC Entry:** 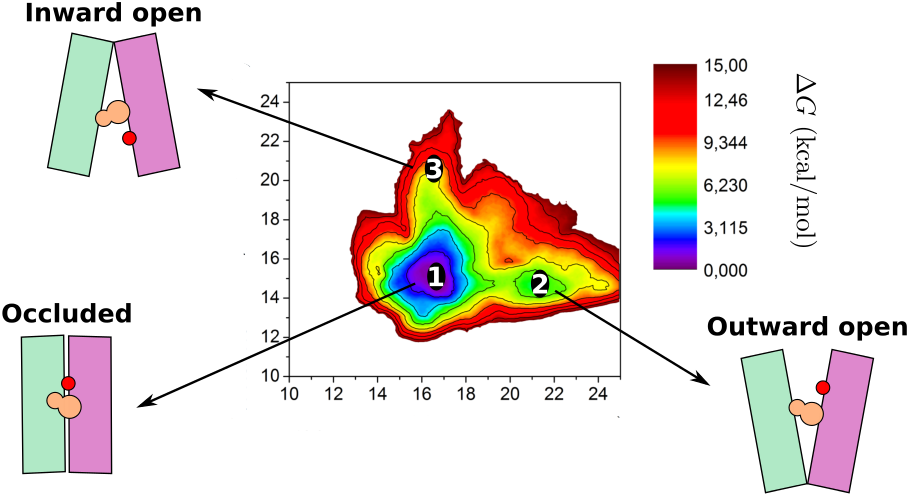

## 1 Introduction

Proton-coupled oligopeptide transporter (POT) is a group of membrane transporters know for their role in the cellular uptake of di- and tripeptides. The mechanism of peptide transport is conserved across all organisms and is one of the main route through which cells absorb nitrogen from the environment.^1,2^ POTs are symporter members of the major facilitator superfamily (MFS) of transporters, which use the inwardly direct proton electrochemical gradient to promote the transport of peptide and peptide-like molecules across the cell membrane. In humans, the POT family transporters, PepT1 (SLC15A1) and PepT2 (SLC15A2), are found predominantly in the small intestine and in the kidney, respectively. Both transporters recognize a diverse library of peptide substrates ^3,4^. In addition to peptides, they also play an active role in the oral absorption and renal retention spectrum of drug compounds that exhibit a steric resemblance to peptides, including *β*-lactam antibiotics and peptiditic prodrugs,^5,6^ which make them some of the most promiscuous transporters. Therefore, the substrate multispecificity of these proteins is of interest in various fields of science, particularly due to its potential relevance to medical applications. However, a detailed mechanism by which these proteins recognize and transport a huge diversity of substrates is currently absent.

The members of POT family exhibit a high degree of sequence conservation. This conservation between prokaryotic and eukaryotic PTR family members highlights the remarkable degree of the mechanistic conservation within this family of transporters. Although mammalian POTs are yet to be crystallized, several bacterial members have been crystallized and their 3D structure determined, ^7–13^ The first POT structure was obtained by Newstead et al in 2011 from the bacterium *Shewanella oneidensis*, PepT_*so*_.^7^ The structures of PepT_*st*_, from *Streptococcus thermophilus*,^8^ GkPOT, from *Geobacillus kaustophilus*^9^ and PepT_*so*2_ also from *S. oneidensis*^14^ followed. Posteriorly, the crystal structure of di- and tripeptide bound complexes of PepT_*st*_ were also determined.^11^ All structures revealed a fold shared by MFS members, consisting of 12 transmembrane (TM) helices, arranged in two bundles (Figure 1A-B). The N- and C-terminal bundles, formed by TM helices H1-H6 and H7-H12, come together to form a “V’ shaped transporter. Bacterial POT members structures have two additional TM helices, named HA and HB, which connect the N- and C-terminal bundles. Although the role of these additional helices is still unclear, their absence in other protein sequences suggests they are not involved in the conserved transport mechanism. ^7,15^ The peptide binding site is a hydrophilic cavity, located at the center of the membrane. This central cavity is formed by residues from H1, H2, H4 and H5 from the N-terminal bundle and from H7, H8, H10 and H11 from the C-terminal bundle. All the crystal structures determined exhibit the extracellular side of the binding site sealed, through the close packing of helices H1 and H2 against H7 and H8, forming an extracellular gate. On the other hand, the access to the binding site from the cytoplasm is restricted by an intracellular gate formed by helices H4 and H5 (N-terminal side) and H10 and H11 (C-terminal side). The interaction between these helices occurs through residues that are conserved across POT members.

**Figure 1:**
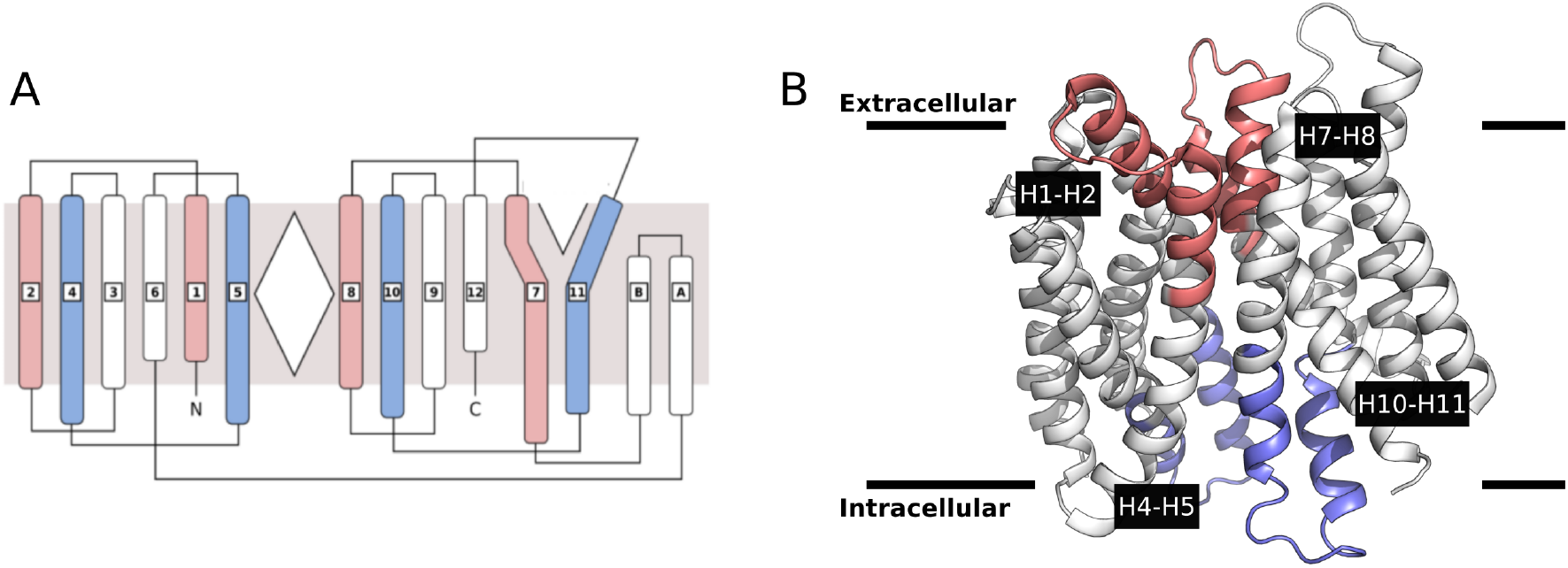
(A) Topology and (B) crystallographic structure of PepT_*st*_. The helices belong to intracellular and extracellular gates are colored in blue and red, respectively.

In order to transport di- and tripeptides across the membrane, membrane transporters must undergo global conformational changes to alternate between inward-facing (IF), outward-facing (OF) and occluded (OC) states, enabling the central peptide binding site to be alternately exposed to either side of the membrane (Figure 2). This model is commonly named as “alternate access mechanism” and describes the transport mechanism of MFS transporters. A “rocker-switch” mechanism, an extension of the alternate access model, was proposed by Solcan et al. for POTs.^8^ According to this model, the alternate access is achieved through rocking of the N-domain and C-domain over a rotation axis that crosses the central substrate binding site at the domain interface. The rocker-switch mechanism can be thought as a highly concerted global conformational change, where N- and C-terminal bundles rotate respective to one another.

**Figure 2:**
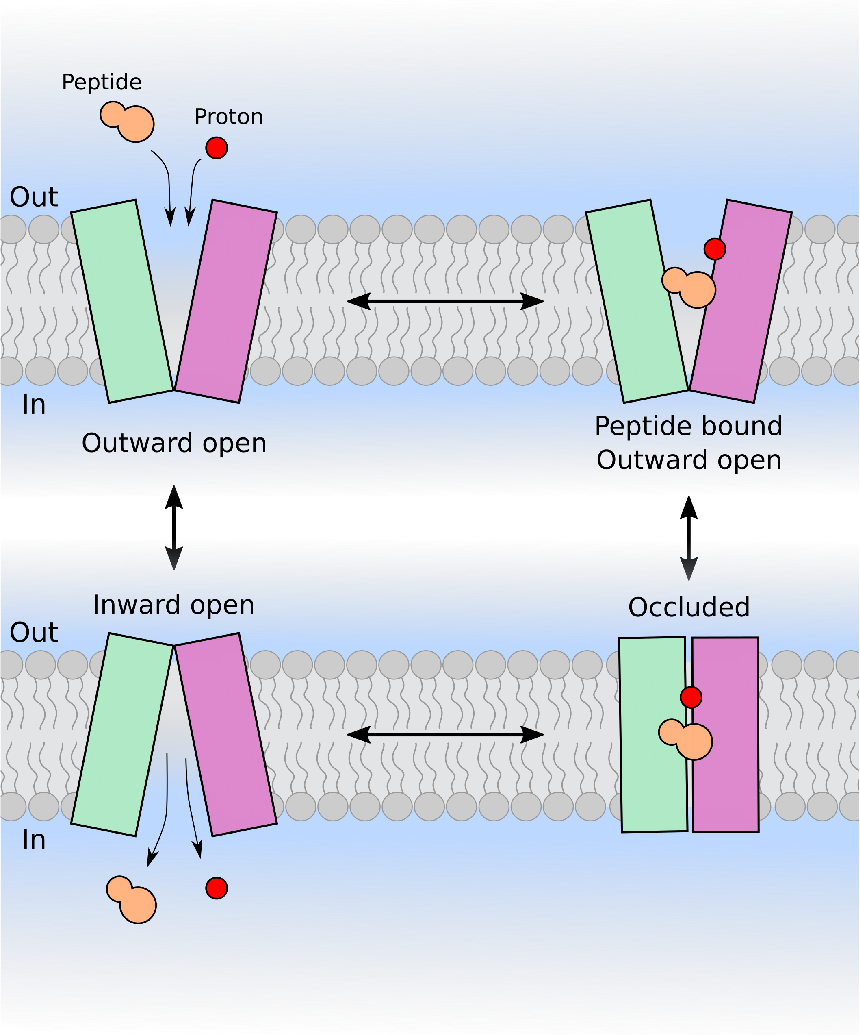
Schematic representation of POT alternate access transport mechanism.

While the structural differences observed in crystallographic models provide information about the structural basis for the mechanism of POT transporters, many mechanistic details remain uncharacterized, partly due to the absence of outward-facing experimental models. All POT crystal structures have been described as inward-facing or partly occluded inward-facing, and thus can not be used to elucidate the IF ↔ OF conformational transitions. Molecular dynamics (MD) simulations have been extensively used to provide a detailed picture of the dynamic of POT proteins^9,10,16–18^ and other membrane transporters.^19–24^ However, the typical time scale of atomistic molecular dynamic simulations is much smaller than that associated with global conformational changes of membrane transporter. Thus, unbiased atomistic simulations is often limited to local conformational dynamics of individual states. Although consistent whit the rocker-switch mechanism, the majority of recently reported MD simulations of POTs have not captured the OF conformation. Parker et al ^17^ have demonstrated that the ensemble of conformations adopted by His67 (TM2) protonated PepT_*Xc*_ is similar to the conformations observed in YajR (PDB ID code 3WDO ^25^) and FucP (PDD ID code 307P^26^) crystallographic structures, in their outward-open states. Nevertheless, the mechanism for proton coupling within POT family can be divided in two types: one that contains the conserved histidine in TM2 and another without the histidine, which includes PepT_*st*_. Using exhaustive MD simulations of GkPOT carried out under 16 different conditions (in terms of the protonation state and presence/absence of substrates) Immadisetty et al. have demonstrated that time scales accessible to typical molecular dynamics simulations can characterize distinct behavior of local conformational dynamics depending on the simulation conditions. However, these local perturbations does not result in any statistically significant differences in the global conformational dynamics of the protein.^16^

In this study, we aim to describe in details the conformational equilibrium of PepT_*st*_ under different simulation conditions using enhanced sampling methodologies. We have conducted an extensive computational study of intermediary states of the transport cycle of PepT_*st*_ in order to describe how the protonation state and the presence/absence of substrate control local and global conformational transitions using an energetic perspective. Our results describe the OF state and provide a view of the large-scale transitions undergone by the cyto and periplasmic gates.

## 2 Materials and Methods

Molecular Dynamics simulations were performed for the PepT_*st*_ starting from the crystallographic models 4APS^8^ and 4D2C,^11^ for the representation of apo and holo transporter, respectively. The first crystallographic structure was used for the construction of the models for: 1) apo-PepT_*st*_ (UP:Apo) and 2) Glu300 protonated apo-PepT_*st*_ (P:Apo). The second structure was used for the simulations of holo-PepT_*st*_ (UP:Holo) and Glu300 protonated holo-PepT_*st*_ (P:Holo). The conformations of the missing loops in the crystallographic structures were generated with Modeller.^27^

Using the CHARMM-GUI web interface,^28,29^ each structure was embedded in a fully hydrated palmitoyl-oleyl-phosphatidylcholine (POPC) bilayer consisting of 314 lipids units and around 27,000 water molecules. K^+^Cl^−^ were explicitly added in a concentration of 150 mM to render the systems neutral. Simulations were performed with CHARMM36 ^30^ force-field for protein and lipids. The TIP3P model was used for water. ^31^ All simulations were carried out with NAMD package. ^32^ The long-range electrostatic interactions were computed with the PME method. ^33^ For Van der Waals interactions, a cutoff of 12 Å was used. The simulations were performed in the isothermal-isobaric ensemble at T=303.15 K and 1 atm using a time-step of 2 fs. Temperature was controlled using Langevin dynamics with a damping coefficient of 10 ps^−1^. The Nosé-Hoover algorithm was used for pressure control, with a piston oscilation period of 200 fs and decay rate of 100 fs. Covalent bonds involving hydrogen atoms in the protein and lipids were constrained to their equilibrium distances using the SHAKE algorithm, while the SETTLE^34^ algorithm was used for water. VMD^35^ was used for visualization and figure preparation.

### 2.1 Adaptive Biasing Force simulations

The simulations performed here aimed the comprehensive profiling of the free-energies involved in conformational transitions of the PepT_*st*_ transporter under different conditions. Overcoming the high barriers that separate the thermodynamic states of interest can be highly inefficient when using conventional Boltzmann weighed sampling. To circumvent the inherent difficulties of Boltzmann sampling, the Adaptive Biasing Force (ABF) method ^36,37^ was used to map the free-energies along reaction coordinates involving the opening and closing of the intracellular and extracellular gates. This methodology is currently one of the most efficient strategies for accelerated conformational sampling in MD simulations.

ABF is a strategy for accelerated sampling and free-energy profiling along a reaction coordinate. The idea behind the adaptive biasing force algorithm is to preserve the dynamic characteristics of the system, while flattening the potential of mean force (PMF) in order to eliminate free-energy barriers and, consequently, accelerating transitions between states. In practice, the instantaneous force acting on atoms that define the reaction coordinate is calculated and averaged. After a good estimation of the average force, an external biasing force, with the same modulus but opposite direction is applied to the system. Then, the total force acting along the reaction coordinate will be on average null, and the system will experience a nearly flat potential of mean force. More details of the method with rigorous descriptions can be obtained elsewhere^36–40^

The ABF simulations were performed as implemented in NAMD. As we aimed to explore the conformational variability associated with transitions between inward-facing, occluded and outward-facing conformations, the reaction coordinate should characterize the movement of opening/closing of the intra- and extracellular gates. A two dimensional free energy landscape was generated using as reaction coordinates two distances: 1) the center of mass (COM) distance between the tips of H1-H2 and H7-H8 and 2) the COM distance between the tips of H4-H5 and H10-H11 (Figure 1A). The tip of an helix was defined as the first (or last) ten residues of the helix. Only the C_*α*_ atoms were considered for calculation. The distances were sampled by ABF simulations within 10 and 25 Å with a precision of 0.1 Å. To restrain the movements of the atoms inside the region of interest, harmonic boundary potential with a 10 kcal mol^−1^Å^2^ force constant was applied. The ABF force was only applied after 5,000 samples of the mean force were generated in each bin. For each system, a total of 2.0 *μ*s of ABF simulations were performed. These simulations were divided in 20 independent sets of 100 ns each, starting from the crystallographic models and from intermediary transport states obtained during the simulations. The convergence of the free-energy profiles was checked by computing the root mean square of the ABF forces. The profiles were considered converged if their root mean square was smaller than 0.2 kcal mol^−1^. For regions with poor sampling, additional windows were inserted to enhance sampling.

## 3 Results and discussion

### 3.1 Accessible Conformations of PepT_*st*_

A definitive way to determine the accessible conformations of a system is through their conformational energy profile. The energy landscape of a protein is generally described as a rough surface, which means that a protein will have multiple energy minima, depending on its environment and interactions. The depth of the energy minima describes the thermodynamic stability of a protein; on the other hand, the heights of the barriers separating energy minima correlate with protein’s kinetic stability, i.e., how readily it can leave one conformation and sample another. Thus, the ability of the protein to sample alternate conformations and the probability with which this sampling occurs is described by their energy landscape. ^41^

The conformational free energy profiles of unprotonated Apo-PepT_*st*_, Glu300 protonated Apo-PepT_*st*_, unprotonated Holo-PepT_*st*_ and Glu300 protonated Holo-PepT_*st*_ are shown in figure 3. The first noticeable difference between the free energy surfaces is that in the protonated models there is a larger region of low free energy. This difference is more evident in the presence of the Ala-Phe dipeptide, where two discernible minimima are present. The narrower free energy well observed for the unprotonated systems indicates that, in this condition, only small perturbations around the global minimum are thermodynamically allowed. After proton binding, the enlargement of the region of low free energy suggests a greater conformational variability relative to the unprotonated systems, and transitions toward OF conformations are then observed. The presence of the second free energy minimum in the Glu300 protonated Holo-PepT_*st*_ system suggests that conformations displaying the periplasmic gate in an open position are not only accessible but also stable. Another distinctive behavior can be observed after Ala-Phe binding, particularly in the position of the global minimum: the position of the global free energy minimum is shifted to regions with lower values of H4-H5/H10-H11 COM distance (y coordinate), i.e., to regions associated with structures exhibiting the closed cytoplasmic gate.

**Figure 3:**
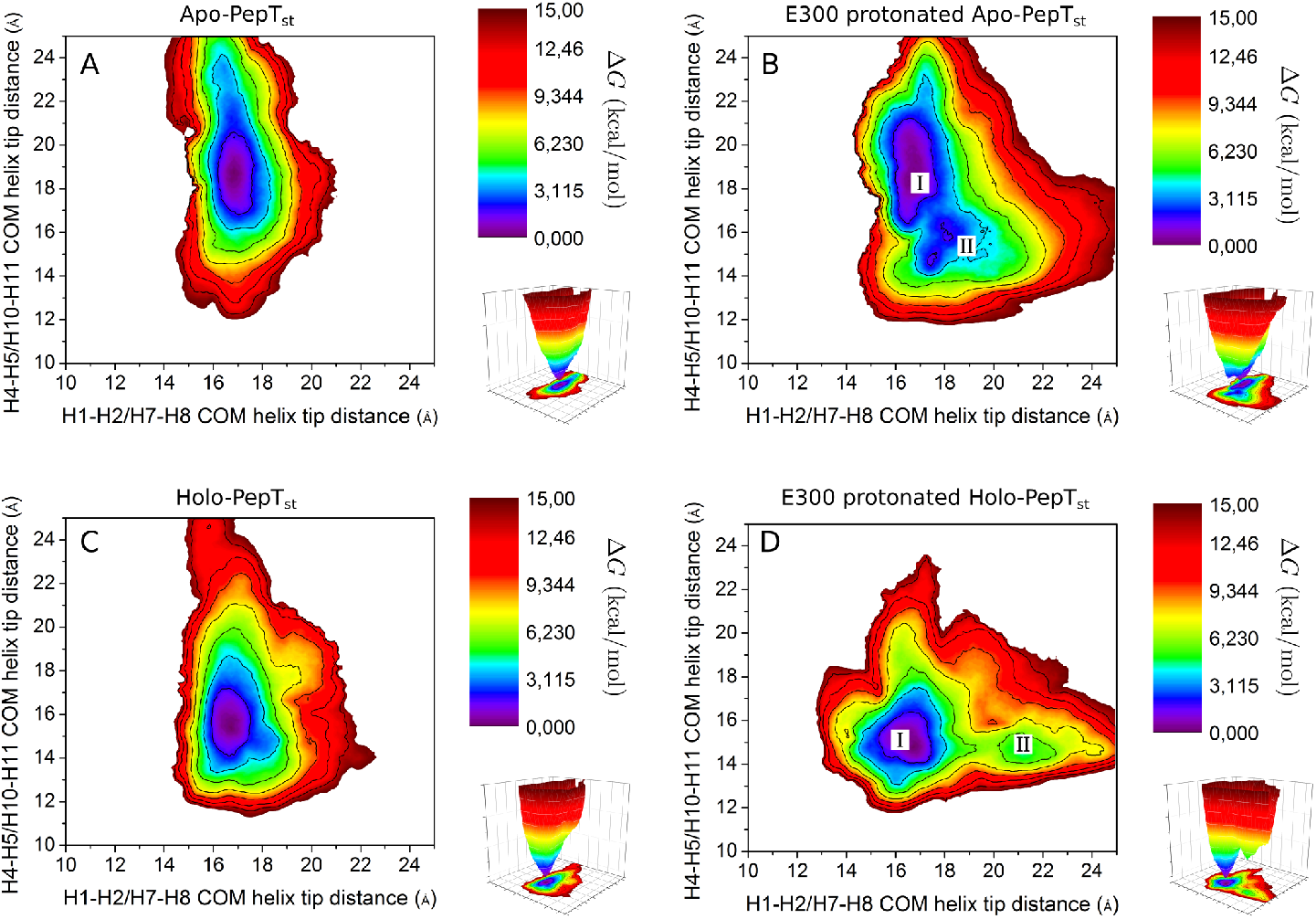
Free energy surfaces of the conformational variability of the (A) Unprotonated Apo-PepT_*st*_, (B) Glu300 Protonated Apo-PepT_*st*_, (C) Unprotonated Holo-PepT_*st*_ and (D) Glu300 Protonated Holo-PepT_*st*_.

The differences in the free energies surfaces allow us to propose general models for proton and ligand related conformational changes of the transporter. The protonation does not directly affect the configuration of the helices that form the cytoplasmic gate, whereas, it facilitates transitions involving OF conformations. The same effect promoted by protonation was observed for PepT_*Xc*_.^17^ While the protonation is related to the conformation of the periplasmic gate, ligand binding seems to perturb the positions of the intracellular gate. The association with the dipeptide Ala-Phe has a destabilizing effect on the IF conformation and promotes the closure of the cytoplasmic gate. Although the OF state is accessible after protonation, its free energy is ~3.5 kcal mol^−1^ higher than that of the global minimum. Therefore, for all systems the most stable states correspond to inward-facing or occluded conformations. This observation is consistent with the fact that PepT_*st*_ crystallizes favorably in one of those states. ^8,11,42^

To characterize the accessible conformations, structures differing in energy by less than ~ 4 kcal mol^−1^ from the global minimum were selected. At 298.15 K, a difference of 4 kcal mol^−1^ represents an upper limit to the observable states. Thus, states differing in energy in more than ~ 4 kcal mol^−1^ are not significantly populated at room temperature. The conformational variability of PepT_*st*_ in all conditions was studied by clustering the accessible structures sampled by ABF simulations using Cpptraj from AmberTools. ^43^ Pairwise RMSDs were used as similarity measure. For the unprotonated systems (Figures 3A and C), two groups of conformations were obtained: one containing structures similar to the occluded conformation, and other composed by structures displaying inward-facing conformation. In the absence of ligand the most populated cluster is the one represented by IF conformations, while for the system bound to Ala-Phe, occluded conformations correspond to the most frequent group of structures. Therefore, without protonation, OF conformations are not observed. After protonation, a new set of conformations was obtained. For the protonated Apo (P:Apo) protein, the accessible structures can be separated in three clusters: the most frequent comprising structures with IF conformation, a second cluster with structures exhibiting occluded conformations and a third minor set of structures containing OF conformations. Finally, for the protonated holo (P:Holo) protein, the accessible conformation are clustered in only two sets of conformations, one represented by the OC state and the other by the OF state.

### 3.2 Conformational transitions induced by protonation and ligand binding

We have used a number of parameters associated with both global and local conformational features of the system to characterize the behavior of the states sampled under different simulation conditions. In order to quantify the global conformational changes we first measured the relative orientation of the N- and C-bundle domains of PepT_*st*_ using the angle between the principal axes of each domain. This quantity is useful to inspect the global conformational changes in POT transporter since it is the main mode of motion, according to the rocker switch mechanism.^8,16^ Figure 4 shows the distribution of the angle between the two domains and the conformation adopted by the representative structure of the global free energy minima under each condition.

**Figure 4:**
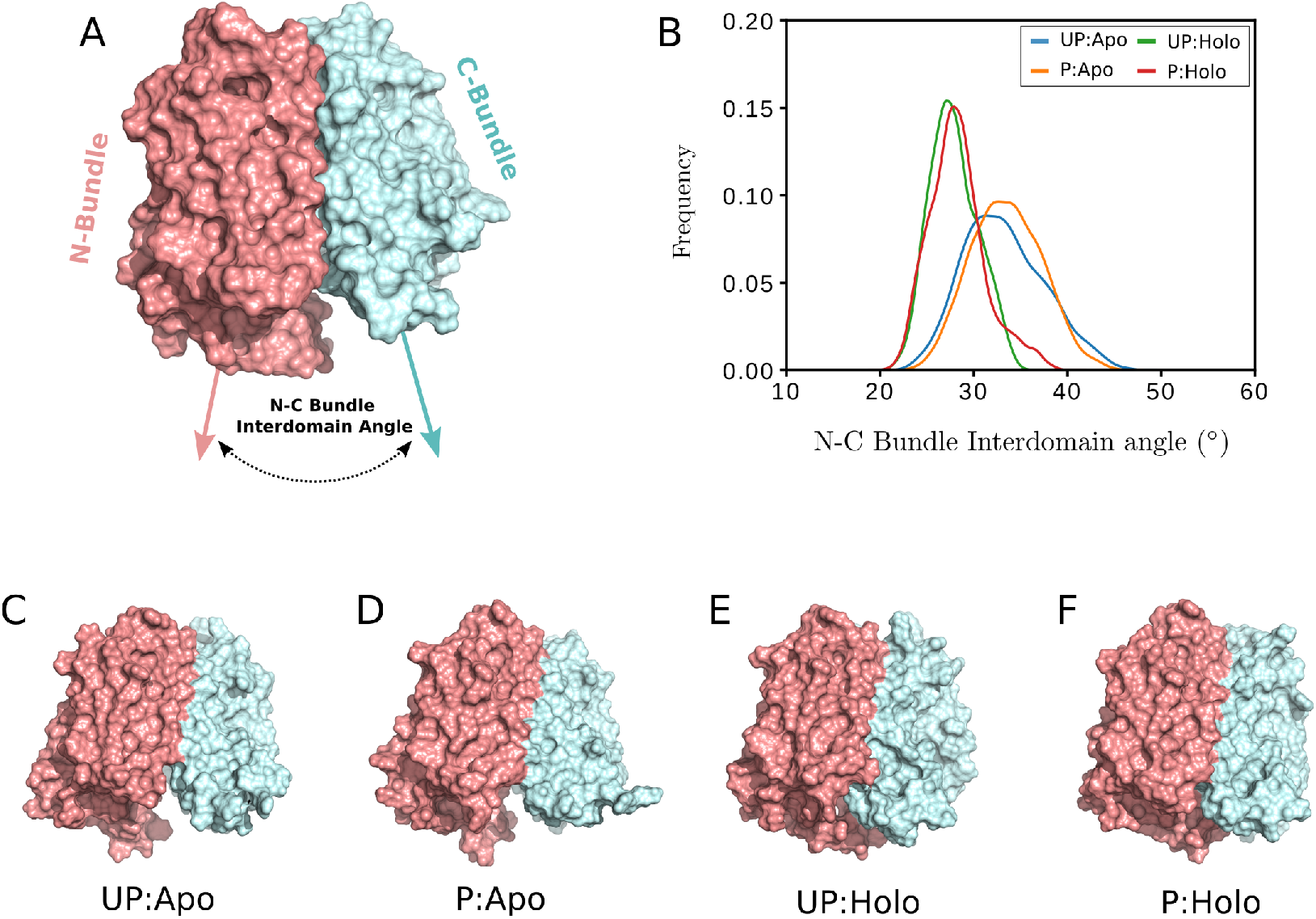
(A) Definition of the N-C bundle interdomain angle. The principal axes of each domain were used. (B) Distribution of the relative orientation of the N- and C-bundle domains for each system. Representative conformation of (C) UP:Apo, (D) P:Apo, (E) UP:Holo e (F) P:Holo, illustrating the relative orientation of N- and C-bundles.

The differences between apo and holo systems are evident. While the distribution for UP:Apo and P:Apo are broad and centered at higher values, UP:Holo and P:Holo systems exhibit narrow distributions. As illustrated in Figures 4C-F, in the absence of the dipeptide Ala-Phe, the cytoplasmic part of the domains are separated from each other, exposing the binding site. The presence of the ligand promotes a concerted motion of the N-bundle and C-bundle, occluding the binding site from the inner side of the membrane. Despite the notable differences, this quantity is not able to distinguish between occluded and outward-facing conformations. Performing the same calculation, but using only the structures located in the second free energy minimum of P:Holo PepT_*st*_ (region II of Figure 3D), i.e, only structures with OF conformation, we obtained a distribution curve similar to that with all accessible structures.

To track the global conformational changes, we also monitored the behavior of the TM helices that form the periplasmic and cytoplasmic gates. As described by Fowler et al, the minimum C_*α*_-C_*α*_ distance between the tips of H4 & H5 and H10 & H11 (L_4,5_-L_10,11_, Figure 5A) and the minimum C_*α*_-C_*α*_ distance between the tips of H1 & H2 and H7 & H8 (L_1,2_-L_7,8_, Figure 5B) correlate with the states of the cytoplasmic and periplasmic gates, respectively. ^10^ The comparison of the distance distributions obtained from the accessible conformations for each system provides some hints about the effect of protonation and ligand on the periplasmic and cytoplasmic gates. In agreement with free energy profiles, the position and widths of L_4,5_-L_10,11_ distance distributions for UP:Apo and P:Apo reflect the greater flexibility of cytoplasmic gate relative to that observed in the presence of Ala-Phe. Therefore, while in the absence of ligand the cytoplasmic gate can alternate between open and closed conformations, for holo-PepT_*st*_ only structures with the cytoplasmic gate adopting a closed conformation are sampled. Monitoring the effect of the interaction with ligand on L_1,2_-L_7,8_ distance distribution indicates that the presence of the dipeptide is not the major aspect interfering on the conformations adopted by the periplasmic gate. However, protonation significantly changes the distance distribution related to the periplasmic gate. The broad distributions observed after protonation suggest an increase in the flexibility of the periplasmic gate and therefore proton binding may represent a trigger to initiate transitions in the direction of OF conformations.

**Figure 5:**
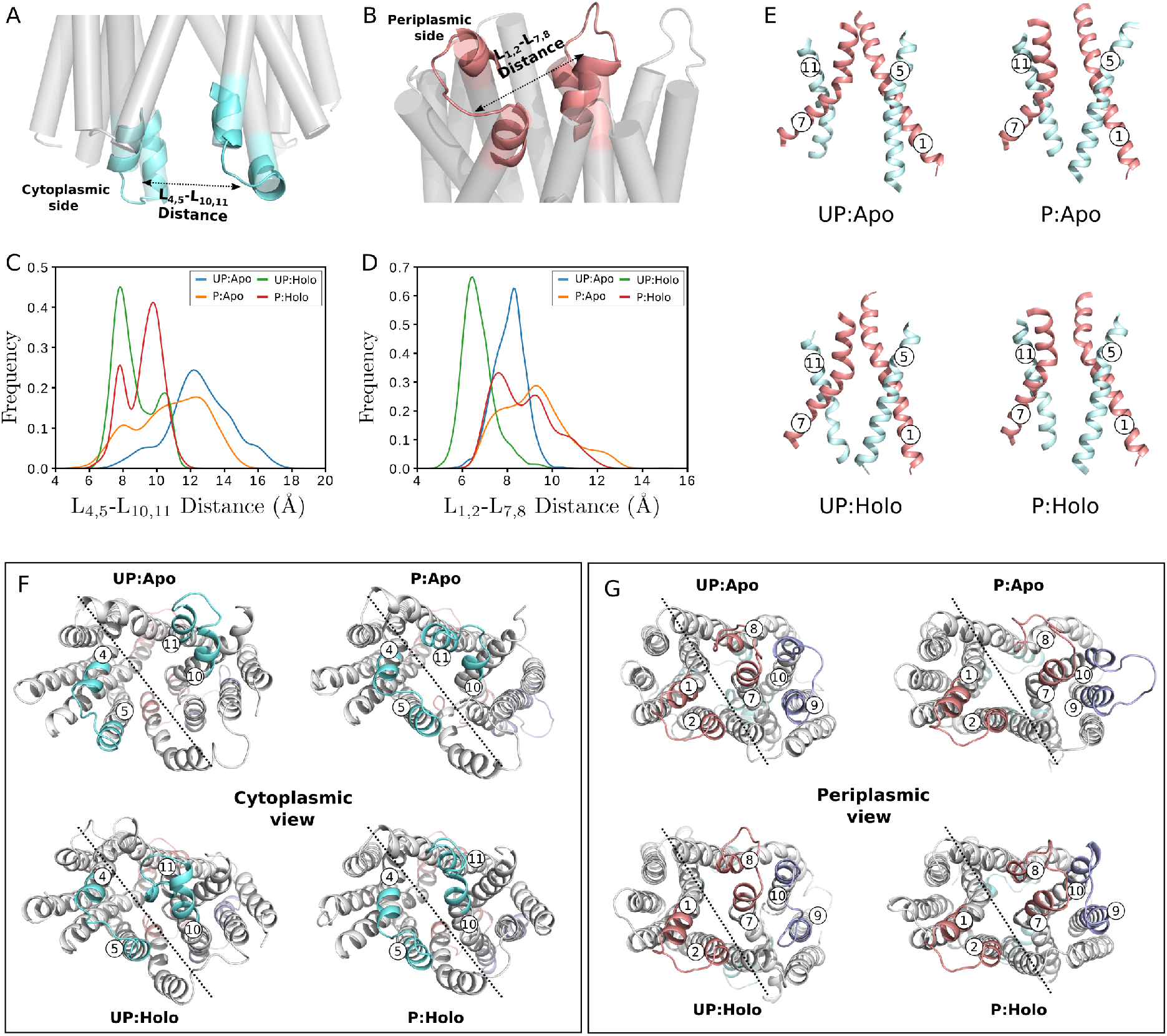
Definition of (A) L_4,5_-L_10,11_ and (B) L_1,2_-L_7,8_ interloop distances used to monitor the opening/closing of the cytoplasmic and periplasmic gate. L_4,5_-L_10,11_ (C) and L_1,2_-L_7,8_ (D) interloop distance distributions. (E) Conformations adopted by helices H5-H11 and H1-H7 for UP:Apo (IF), P:Apo (OF), UP:Holo (OC) and P:Holo (OF). Conformational changes of periplasmic (F) and cytoplasmic gates (G) of PepT_*st*_: The dashed lines approximately represents the plane of symmetry. TM helices of cytoplasmic (H4-H5 and H10-H11) and periplasmic (H1-H2 and H7-H8) are colored in cyan and red, respectively. Helices H9-H10 are colored in blue. For unprotonated systems the conformations of the global minima are represented, while for protonated systems, the most stable conformations of region II (Figure 3) are shown, i.e. the occluded state. The representative conformations of global free energy minima for P:Apo and P:Holo are very similar to that observed for UP:Apo and UP:Holo, respectively

In addition, interhelical angles were also monitored, to capture the movements of the periplasmic and cytoplasmic gates. These angles were used to describe conformational changes on GkPOT^16^ and other membrane transporters.^23,44^ To measure the cytoplasmic gate mobility the angle between the roll axes of helices H5 (N-bundle) and H11 (C-Bundle) were calculated. Likewise, the behavior of the periplasmic gate was captured by monitoring the angle between helices H1 (N-bundle) and H7 (C-Bundle). These quantities may not be able to characterize the different conformations that membrane transporters can assume since the interhelical angles are defined based on the roll axes of the helices and, therefore, they are considered rigid. However, as illustrated in Figure 5E the helices are flexible and have a complex nonlinear behavior. The opening of the periplasmic gate is associated with the bending of helix 7, while the kinking of helix 11 promotes the opening of the cytoplasmic gate. Similar behavior was previously observed.^8,10^ Monitoring the motions of the helices that form the peri- and cytoplasmic gates by the interloop distances and the interhelical angles provides an effective representation of the global conformational changes induced by protonation and ligand binding. Figures 5F-G show the rearrangements of these helices involved in IF↔OF transitions. The opening of the periplasmic gate is also accompanied by motions of helices H9 and H10 and the loop connecting these helices. The rearrangement of this part of the protein creates space to accommodate helices H7 and H8 in an open conformation.

Finally, we monitored the water accessibility of both cyto- and periplasmic gates to characterize the details of the alternate access mechanism. This quantity is also an indirect way to measure the substrate accessibility. Figures 6A-B show the distribution of the number of water molecules that can access the lumen from each side of the membrane. In the absence of protonation and ligand, water molecules can access the central cavity only through the cytoplasmic gate. After protonation, there is an increase in the number of water molecules in the periplasmic gate due to the opening of this gate. It is interesting to note that for P:Apo both distributions are large, which implies the coexistence of all the states in the transport process. After ligand binding the number of water molecules inside the cytoplasmic gate decreases substantially. Although for UP:Holo PepT_*st*_ the access of the central cavity is blocked from both sides of the membrane, the presence of two peaks for the P:Holo distribution of the number of water inside the periplasmic gate indicates that both OC and OF states are visited.

**Figure 6:**
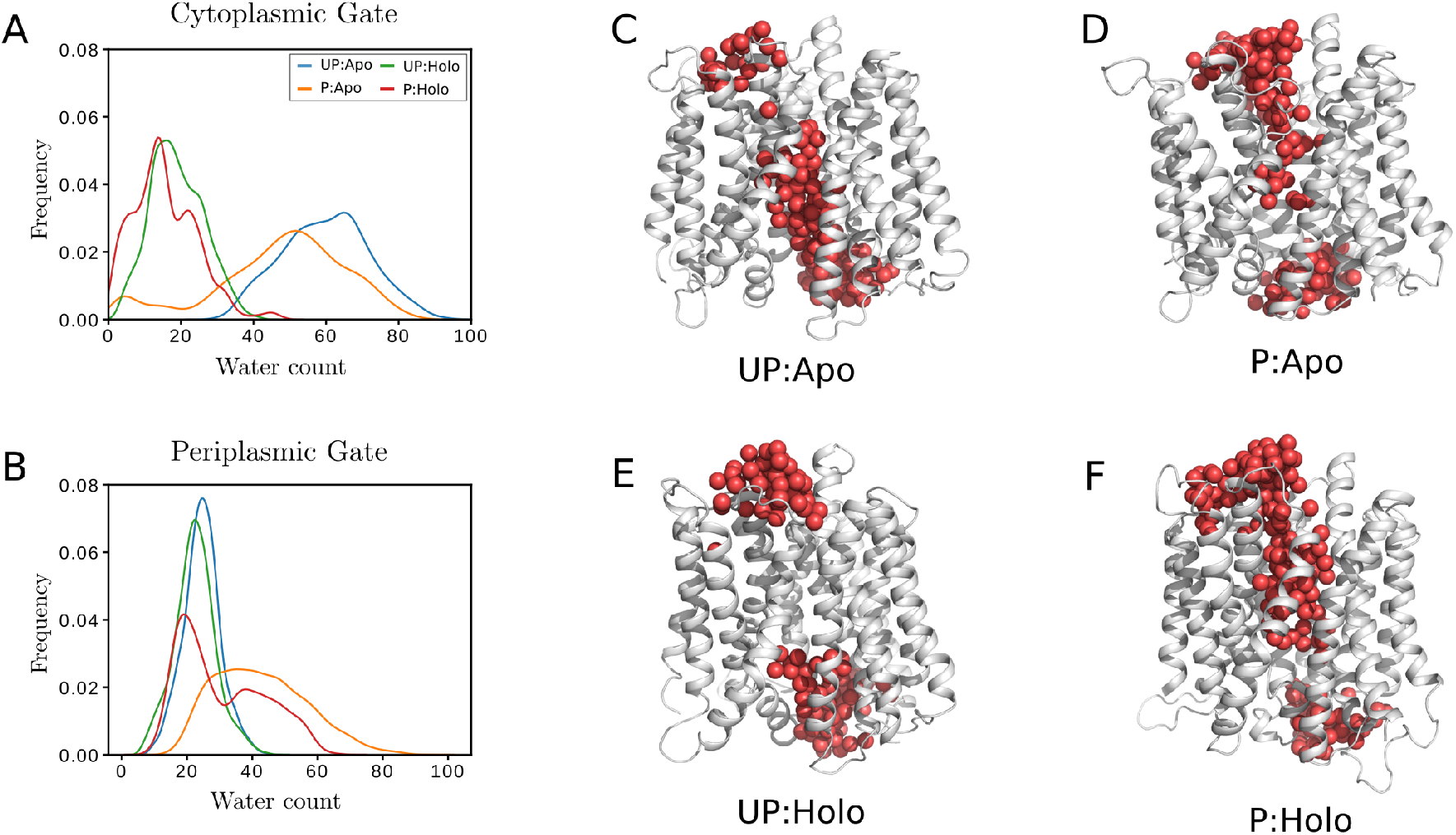
Distribution of water count inside the (A) cytoplasmic and (B) periplasmic gates. Representation of water accessibility for (C) UP:Apo, (D) P:Apo, (E) UP:Holo and (F) P:Holo. Water molecular are represented as red spheres. The same structures shown in Figure 5 were used.

### 3.3 Local conformational changes

The global conformational transitions of PepT_*st*_ are accompanied by local conformational changes, specially within the binding site. To monitor the local dynamics of the binding site, the occurrence of the salt-bridge R33-E300 was calculated. This salt-brigde has been previously suggested to stabilize IF conformations.^8,10^ The equivalent interaction (R43-E310) was also detected in GkPOT transporter.^9^ Combining functional assays with GkPOT mutants and some MD simulations, Doki et al concluded that Arg43 and Glu310 play an important role controlling IF↔OC transitions. However, their hypothesis can not explain the transition mechanism between OF and OC states. In order to track the formation of the R33-E300 salt-brigde, we measured the R33-E300 distance (defined as the minimum distance between the potential hydrogen bond donors and acceptors of R33 and E300 side chains - Figure 7A) for all accessible conformations obtained for each system. Figure 7B clearly illustrates a distinct behavior of the R33-E300 distance distribution in unprotonated and protonated systems. As expected, after protonation, Glu300 cannot form a salt-bridge with Arg33.

**Figure 7:**
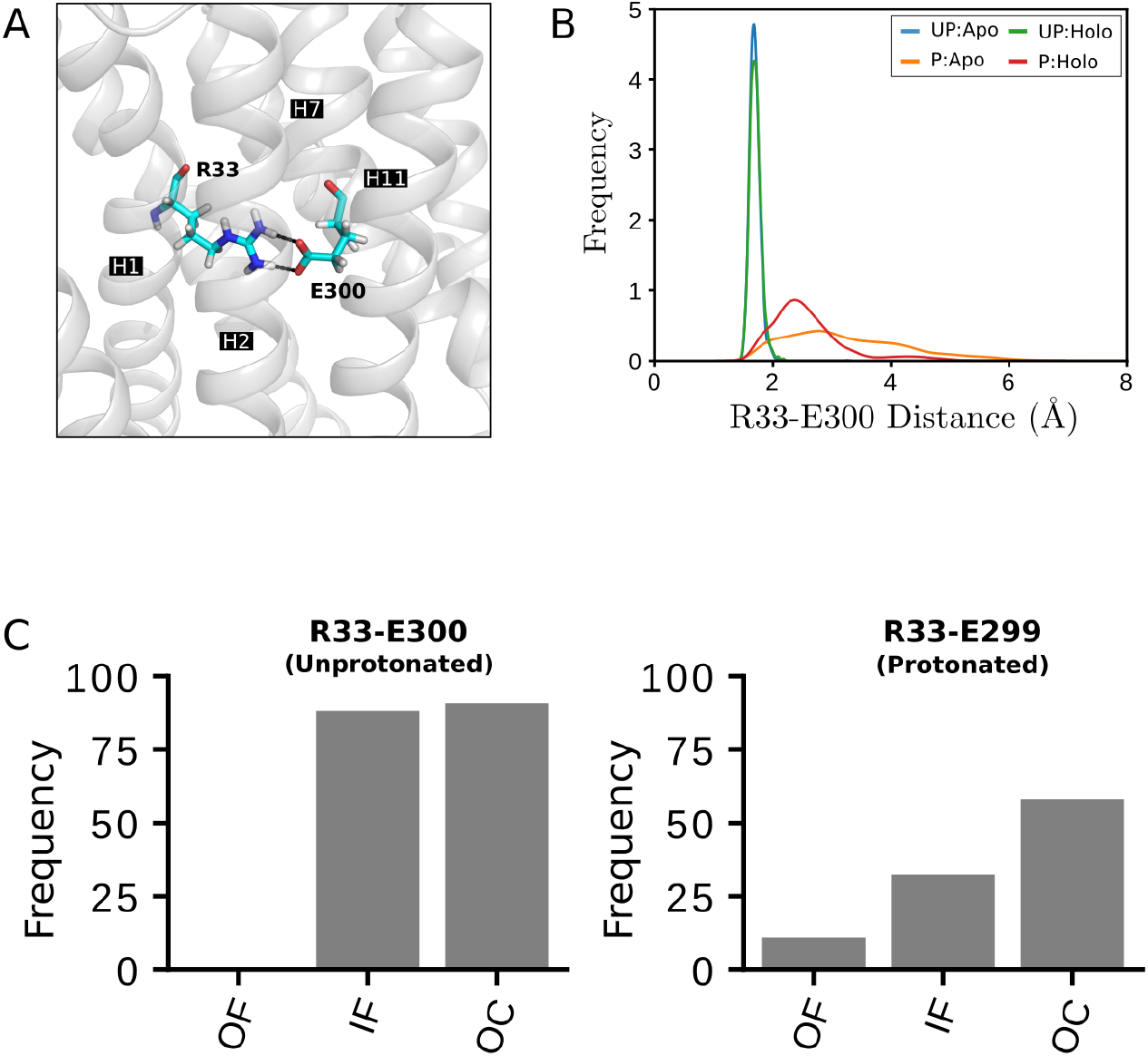
(A) Definition of R33-E300 salt-bridge minimum donor-acceptor distance. (B) Distribution of R33-E300 distance for each simulation condition. (C) Frequency of the formation of the R33-E300 and R33-E299 salt-brigdes for in the different conformations for unprotonated and protonated PepT_*st*_.

The calculation of the frequency of occurrence of the R33-E300 salt-bridge for each state of the transport mechanism reveals that this interaction occurs approximately in 90% of all the structures adopting the occluded and inward facing conformations for unprotonated systems. Therefore, the hypothesis that those salt-bridges play a pivotal role controlling transitions toward outward facing conformations is supported. However, even after protonation, we observed that OC conformations are stable, as shown by the free energy profiles. Analysis of the structures in the OC state for protonated systems identified a new salt-bridge that occurs only after protonation. Protonation disrupts the R33-E300 salt-bridge, however, in protonated systems, this interaction is replaced by R33-E299 salt-bridge. Although this new salt-brigde is less frequent than R33-E300, it has been observed in around 60% of all structures adopting the OC conformation for P:Apo and P:Holo systems and may be important controlling OC↔OF transitions (Figure 7C). Glu299 was proposed as being essential to maintain the structure of PepT_*st*_. Mutations in this residue to alanine, glutamine or conservative aspartic acid resulted in a protein too unstable to be produced. ^8^

Other salt-bridges were identified in the accessible conformations of PepT_*st*_ (Figure 8). Three of these (E22-K126, E25-K126, R53-E312) have been previously described by Fowler et al.^10^ The first two salt-bridges were found to be uniformly distributed across all the states. The third interaction (R53-E312) only occurs in around 15% and *5%* of IF and OC conformations, respectively, and should not be critical to maintain these conformations. Mutations in any of Glu22, Glu25 or Lys126 has been shown to abolish or reduce transport.^8^ Two of the observed salt-bridges (D79-K413 and D145-K347) occur only in structures adopting the OF conformation. They are located at the cytoplasmic side of the membrane and probably play an important role in the stabilization of outward-facing conformations. The residue Asp79 is conserved in members of POT family. Mutating the equivalent residue in PepT_*so*_ to alanine significantly reduces transport, which is consistent with the idea that the D79-K413 salt bridge stabilizes OF conformations.^10^

**Figure 8:**
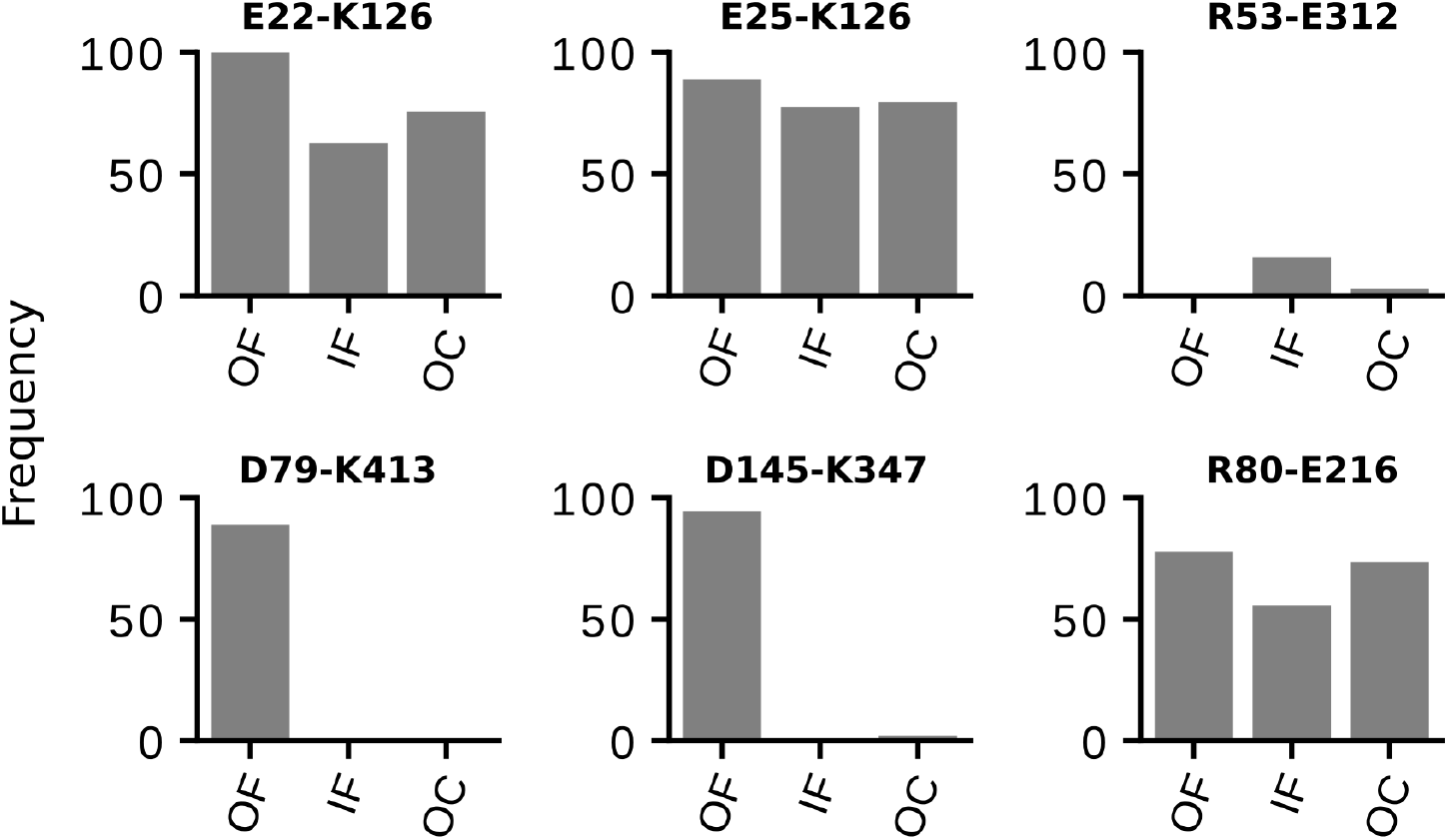
Salt-bridges identified in the different conformations of PepT_*st*_. E22-K126, E25-K126 and R80-E216 was found in all the conformations. R53-E312 salt-brigde occurs in only 15% of the structures adopting the IF conformation and hence is unlike to be critical in the stabilization of this conformation. The last to salt-bridges (D79-K413 and D145-K347) occurs on the cytoplasmic side of PepT_*st*_ and are observed only in the OC conformation.

## 4 Conclusions

We have performed a series of adaptive biasing force molecular dynamics simulations to map the conformational free energy profile of PepT_*st*_ and study how protonation and peptide binding affects the dynamic properties of this transporter.

The free energy profiles obtained allowed us to propose general models for proton and ligand related conformational changes of the transporter. In the absence of ligand and protonation, PepT_*st*_ is able to shit between IF and OC conformations. However, OF conformation are not thermodynamically accessible. In this condition, the most stable conformation is the inward-facing state, justifying the fact that the majority of the POTs crystallographic structures are obtained in this conformation.

Protonation perturbs the free energy surface and the differences in the free energy profiles clearly indicate a distinct behavior in the dynamic of PepT_*st*_. After protonation in Glu300, the Arg33-Glu300 salt-bridge is broken, which increases the flexibility of the periplasmic gate. In the absence of ligand, all the three states (IF, OC and OF) of the transport mechanism are thermodynamically accessible. However, only the inward-facing conformation corresponds to a stable state, i.e., is associated with a free energy minimum. The differences in the accessible conformations before and after protonation suggest that the protonation is essential to promote conformational transitions toward the OF conformation, and this might be the initial step in the mechanism of transport.

Ligand binding, on the other side, does not seem to perturb the periplasmic gate. The narrow free energy well obtained indicates a high stabilization of the occluded conformation, which is supported by all the measures associated with global and local conformational changes. In the presence of ligand and protonation, two minima with small free are present, and transitions between these minima are also of low energy. The presence of the second free energy minimum localized in coordinates associated with OF conformation indicates that protonation and ligand binding stabilizes the outward-facing conformation. Two salt-bridges (Asp79-Lys413 and Asp145-Lys347) were observed only in structures exhibiting outward-facing conformation, and were predicted to stabilize this conformation. At the same time, the presence of Ala-Phe after protonation perturbs the inward-facing conformation, increasing its energy.

This study presented a detailed picture of local and global conformational changes in terms of various parameters. Although our simulations confirm the global motions of the the N- and C-terminal domains relative to one another, a literal rocker-switch mechanism, based on rigid domain movements, is not sufficient to describe IF↔OF transitions. Our results indicate the coupling between local conformational changes and the flexibility of cyto- and periplasmic gates. In conjunction with previous experimental and simulation data, this work contributes toward a definitive model of POTs transport mechanism.

## Acknowledgement

The authors thank Prof. Simon Newsted and Dr. Joanne Parker (Biochemistry Department - University of Oxford) for valuable discussions and Fundação de Amparo à Pesquisa do Estado de São Paulo (Grants 2015/50366-7 and 2016/16328-3) for financial support.

